# A Generative Model for Creating Path Delineated Helical Proteins

**DOI:** 10.1101/2023.05.24.542095

**Authors:** Ryan D. Kibler, Basile Wicky, Brian Coventry, Nicholas B. Woodall

## Abstract

Engineered proteins with precisely defined shapes can scaffold functional protein domains in 3D space to fine-tune their functions, such as the regulation of cellular signaling by ligand positioning or the design of self-assembling protein materials with specific forms. Methods for simply and efficiently generating the protein backbones to initiate these design processes remain limited. In this work, we develop a lightweight neural network to guide helix fragment assembly along a guideline using a GAN architecture and show that this approach can rapidly generate viable samples while being computationally inexpensive. Key to our approach is the transformation of the input structural data used for training into a parametric representation of helices to reduce the generator network size, which in turn facilitates rapid backpropagation to find specific helical arrangements during generation. This approach provides a method to quickly generate helical protein scaffolds.

## Introduction

Deep learning methods have accelerated all aspects of protein design including backbone generation^1–5^, sequence prediction^6–8^ and design confirmation^9^. Already, the AlphaFold2 network has revolutionized the prediction of protein structure from sequence data alone, illustrating the power of deep learning methods to learn from protein data sets^10^. Designing *de novo* proteins with machine learning methods has found many successes, but fast and accurate backbone generation methods are still limited. Here, we have developed a series of protein design tools using machine learning and parametric methods to efficiently generate four-helix protein backbones and design their sequence. These four-helix units can be combined to create larger single-chain proteins of desired shapes by backpropagating through the neural network to find the right combinations of four-helix folding cores.

Machine learning has contributed several methods to the first step of *de novo* protein design, namely backbone generation. Methods like hallucination modify the sequence passed to structure prediction networks like trRosetta, RoseTTAFold or AlphaFold until convergence to a protein backbone with desired structural features. Leveraging the accuracy of these networks, an entire protein can be designed around a specified structural motif by optimizing a loss function to the design via repeated backpropagation from the structural output to the sequence input of the networks^1,2,6,7,11,12^. Other machine learning networks have more directly generated structures. One used a Generator Adversarial Network (GAN) architecture to make 64-residue protein fragments and another generated immunoglobulin-folds with a Variational Auto-Encoder (VAE)^13–15^. GAN and VAE networks produce proteins that respect the distributions from their learned datasets. However, to achieve a design goal like a specific shape, these approaches still require backpropagation through the network in order to optimize the design loss that will satisfy the characteristics of the desired protein which can be computationally expensive. Recently, generative models relying on diffusion have given promising results towards protein generation, but these models remain computationally expensive^4,5^.

Here we chose to focus on designing helical proteins because they have been reliable protein design targets due to the locally-satisfied backbone hydrogen bonds of the helix. Many recent papers that claim functional *de novo* designed proteins utilize helical proteins with short loops^16–23^. Repeated helical proteins are also often used as a base to iterate on larger structural architectures^24–27^. The generation of these helical backbones is commonly done by randomly or parametrically assembling sets of structural fragments^28–31^. Our goal is to make this fragment assembly more directed by replacing the random/parametric sampling with a guided assembly using a neural network trained on idealized helical proteins.

Towards this aim, we trained a generator to produce a four-helical protein with the goal of keeping the network small to ensure fast and inexpensive backpropagation during design. To achieve this goal, the input structural data was reduced from an all-atom to a parametric representation by fitting the four-helix dataset to a set of helical parameters using ideal straight helices reducing the input features. The output of the generator guides a fragment assembly process which diversifies the structures by the helical phase.

In addition to this backbone generator network, we also re-trained a graph-based message passing neural network that predicts sequences from input backbone to be compatible with the generated straight-helices^7^. Finally, we show that the network can produce larger helical proteins composed of four-helix units by backpropagation for the required unit at each addition.

## RESULTS

### Straight Helical Fits

Our training dataset consists of 27,877 *de novo* designed small 4-helix bundle proteins, each 65 amino acids long. These are idealized proteins consisting of a single hydrophobic core and short loops. The designed sequences promote the desired secondary structure using many proline and glycine residues in the loops (features the Rosetta energy function needs help in designing). These proteins represent a diverse set of four-helix proteins that can be assembled. Building from a previous helix parameterization method^32^, we fit each of the four helices separately as a straight helix (excluding the loop regions). Each helix is represented by seven parameters: three coordinate parameters (x, y, z) to describe the translation of the helix midpoints from the frame of reference, three Euler angle parameters (Φ, Ψ, Θ) to describe the rotation of the helical axis vector and a single helical phase (φ) parameter to represent the rotation of the helix around its central axis (Fig. 1A). The straight helices fit this dataset with high accuracy (median RMSD = 0.4 Å), with the worst helical fit for each set of four helices being below 1.0 Å RMSD of the C_α_ atoms for 96% of the dataset (Fig. 1 B,C). Next, we converted these parameters to a distance map of helical endpoints for a rotational/translational invariant 3D representation for machine learning. This distance map contains 28 features per 4-helix protein (Fig. 2A) and excluded the helical phase. Since the helical phase of each helix depends on the loop that connects the helices, leaving these parameters out of the generator enables downstream diversification during the looping process (see below).

**Figure 1.**
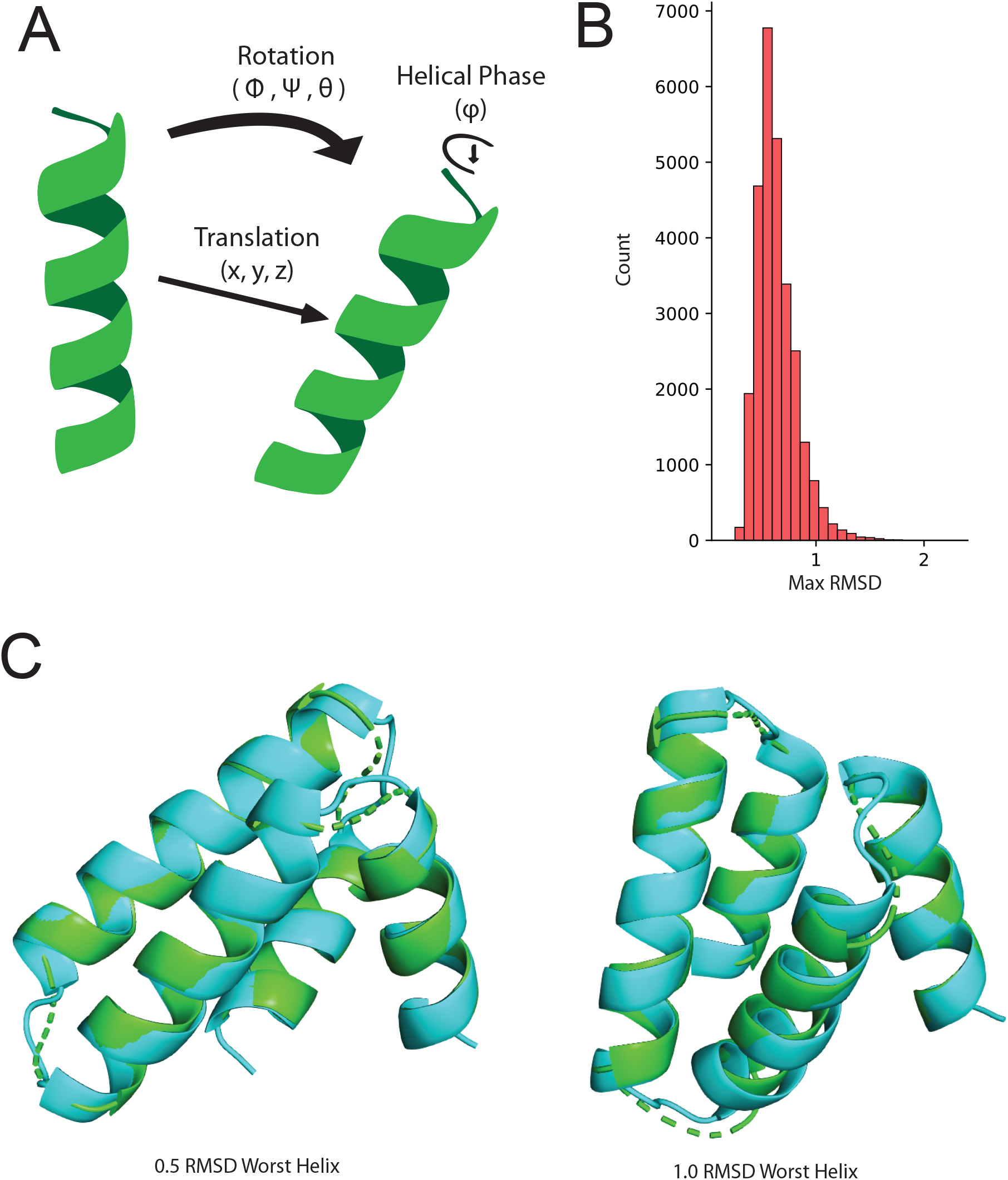
Parametric Helical Fitting **(A)** Diagram of the seven parameters fit to each helix as a straight helix **(B)** Histogram of the RMSD of the worst-fitting helix from each protein in the dataset. **(C)** (Left) Example of a 0.5 angstrom RMSD worst-helix protein. (Right) Example of a 1.0 angstrom worst-helix protein.

**Figure 2.**
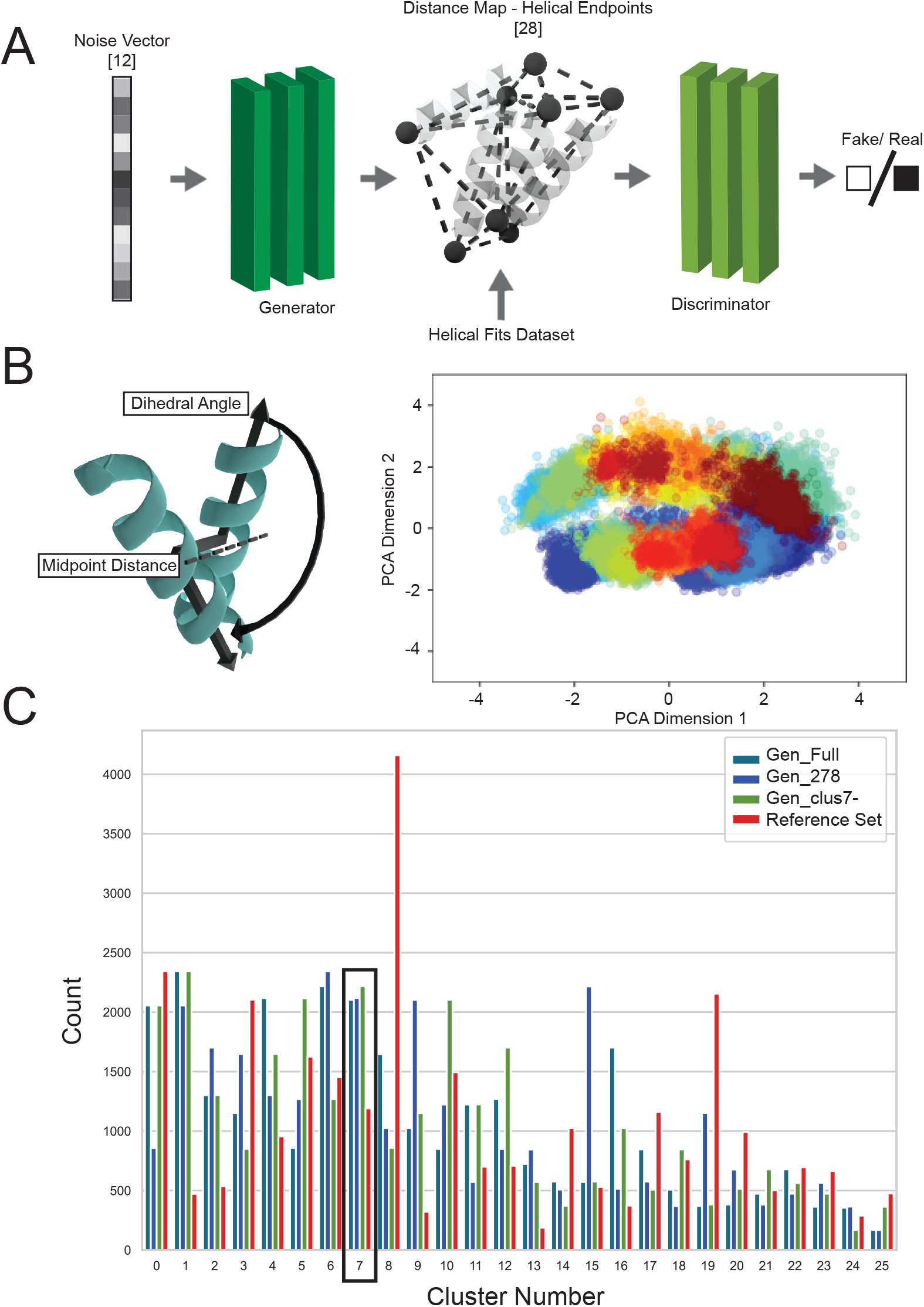
Generator Training **(A)** Diagram of GAN setup **(B)** Parameters used for clustering dataset (left). The first two dimensions of the principal component analysis for the clustering parameters (left). The plot is colored by the clusters. **(C)** Histogram of the clustering of the outputs by the three generators and the reference set. Cluster 7 is highlighted with a black block since Gen_clus7- was trained without cluster 7 but can still produce proteins in cluster 7.

### GAN

A generative adversarial network (GAN) architecture was used for training. The generator is trained to take random vectors and transform them to match the distribution of the training dataset^33^. During training, a discriminator network is trained alongside the generator network to distinguish between the real data and the ‘fake’ output of the generator. Each network iteratively trains the other. As the discriminator becomes better at distinguishing real from fake outputs, it guides the generator. Given the simplification of the helical parameterization, we utilized a small dense neural net of 3 layers by 64 units for both the discriminator and generator (Fig. 2A). We trained the GAN for 500 epochs producing the generator, Gen_Full.

### Generator Evaluation

The output of the generator is a distance map (28 unique features) that is then converted to 8 helical endpoints using Sci-kit Learn’s multidimensional scaling algorithm^34^. Straight poly-alanine helices placed between the endpoints represent the helical arrangement anticipated by the generator assuming a phase of 0°. The ability of the generated backbones to be sequence-designed towards well-packed proteins was evaluated with two backbone (sequence-agnostic) metrics to ensure that helices are close enough to each other to form a hydrophobic core (percent core) while also ensuring the atoms do not sterically clash. For comparison, the reference dataset fits were converted to helical endpoints and then to straight helices as well. There are no steric clashes in the reference dataset but after conversion to straight helical fits 77% had 3 atom clashes or less (5 atoms per residue) (Table 1). The straight fits are approximate and without the helical phase information the directionality of the C_α_/C_β_ vector will differ from the original structure, thus introducing some steric clashes. The generator compared favorably with the reference dataset making 74% of the backbones with less than 3 clashes after generating 27,877 backbones. For the percent ‘hydrophobic’ core metric, based on C_α_/C_β_ atom vector overlap, the generator averaged 13% ± 5% compared to 15% ± 5% for the reference set (Table 1).

**Table 1.**

| Generator Name | gen_full | gen_278 | Reference |
| --- | --- | --- | --- |
| Training Examples | 27877 | 278 | - |
| Reconstruction Error (Å) | 0.12 | 0.12 | - |
| Time (s) per Structure | 0.03 | 0.03 | - |
| Percent No Clashes | 29.5% | 29.1% | 30.2% |
| < 3 Atoms Clashed | 74.7% | 74.6% | 76.9% |
| Clashed Atom Mean | 1.67<br>±1.39 | 1.68<br>±1.30 | 1.61<br>±1.26 |
| Percent Core | 13<br>±5 | 13<br>±5 | 15<br>±5 |

To ensure that the generator creates a diversity of structures instead of a single mode (mode collapse), we first define the diversity of the reference set by structural clustering. The distance map representation was converted to be the helical midpoint distances as well as the dihedral angles of the vectors from the helical midpoint to the C-terminal helical endpoint (Fig. 2B). These 16 features were clustered by spectral clustering into 26 clusters^35^. In order to add new data to the spectral net clusters, we utilized Spectral Neural Net to learn the graph Laplacian transform for the dataset^36^. Newly generated helix backbones can now be clustered and compared to the original reference clusters. The network generated four-helix proteins across all clusters (Fig. 2C). As a further test of the ability of the network to generalize, we arbitrarily removed a cluster, cluster 7, from the reference dataset and trained a new generator network, Gen_clus7-. This new generator was still capable of producing proteins in cluster 7 despite its removal from the training set (Fig. 2C).

One difficulty with training machine learning models on certain classes of protein structures has been the limited number of examples available for training. The reduction in features via the parameterization of the helices (28 features) significantly reduces the number of training examples required to train an appropriately regularized model. As a proof of concept, we trained a new generator based on 278 examples randomly sampled from the reference set, Gen_278. Strikingly, with only 278 training examples, Gen_278 compared favorably to the reference set and Gen_Full in terms of percent core and atom clashing metrics (Table 1).

#### Protein Design Via Network Search

These generators take a vector of random numbers and use them to generate a random protein that falls within the learned distribution. The appeal of *de novo* protein design is to generate a specific protein that solves a specific problem. We show here the utility of the small network size of Gen_Full for backpropagation towards generating specific helical assemblies from four-helix units. As a proof of concept, we generated a helical protein that follows a guide composed of quarter circle that transitions to a line. The underlying principle is to repeatedly search the network for two-helices that match the guide and the previous two helices.

The geometric arrangement of two previous helices can be set as a loss function that is defined as the mean squared error of the desired distance map subtracted by the output distance map with an applied mask to the first two helices (Fig. 3A). This loss function is then minimized by stochastic gradient descent through the network starting from a random input vector to find a new input vector that produces the desired protein.

**Figure 3.**
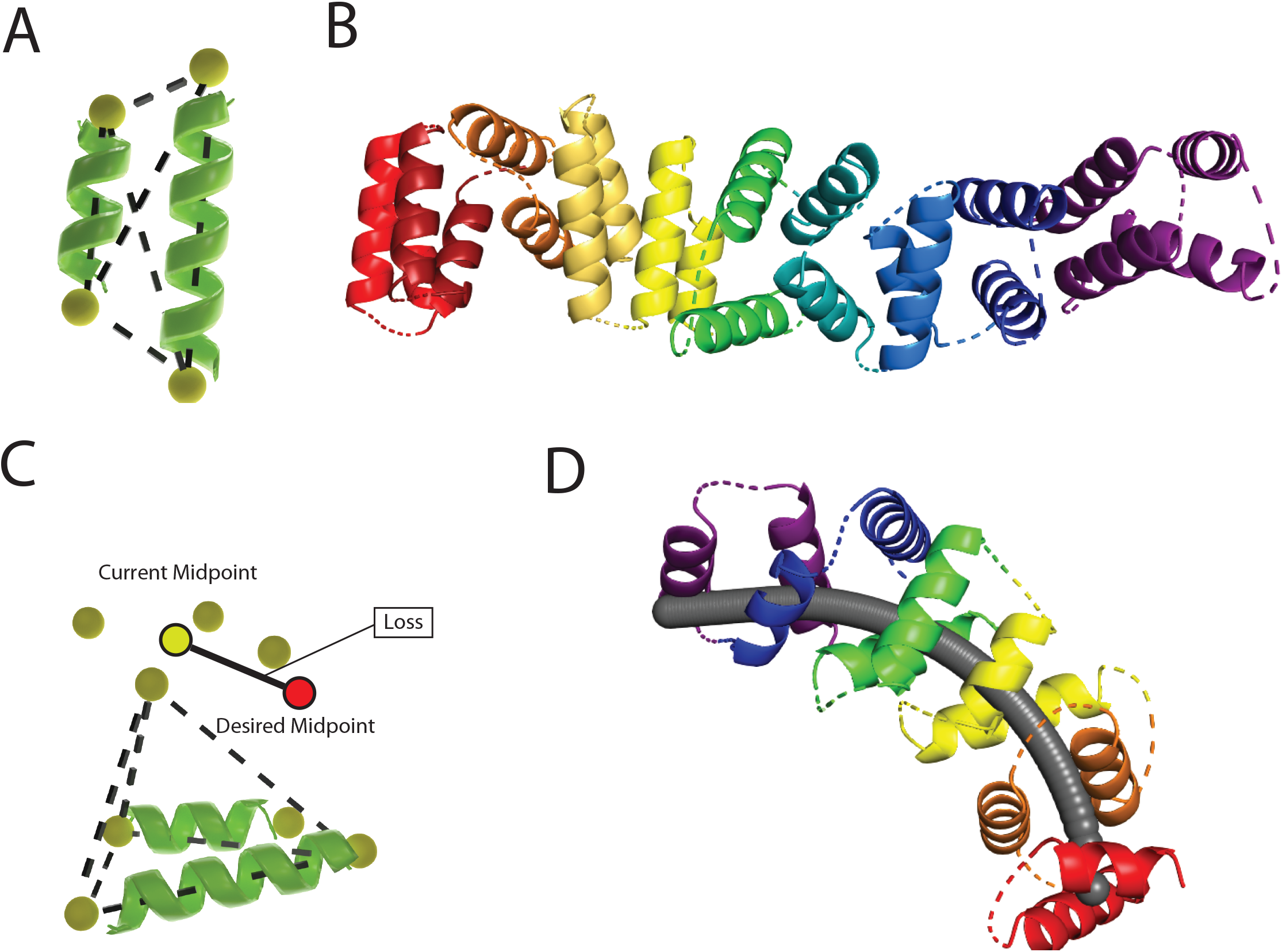
Helical Protein Assembly **(A)** Distances used to map the first two inputs helices into the loss function. **(B)** Example from iterative two helix addition without guides. Straight helices shown are interpolated between endpoints. Dashes show helical connectivity but are not loops. **(C)** Example showing point location calculation from 6 distances from trilateration and midpoints loss. **(D)** One set of endpoints generated along the quarter circle/line guide. Straight helices shown are interpolated between endpoints. Dashes show helical connectivity but are not loops.

Essentially, the network will predict two new helices that will buttress the input two helices. We repeated this process ten times to produce 45 sets of guide endpoints in ∼1.5 minutes on a GPU. An example with straight helices interpolated is shown in Figure 3B. Generally, this process produces helices along a straight line as this is the most probable direction of addition.

Ensuring that the two output helices match the orientation of the guide is more complicated as the generator operates on a distance map. One solution is to convert the distance map of the generator to coordinate space. In a distance map, the minimally determined 3-dimensional shape is a tetrahedron composed of six distances. Solving for a single point can be done by trilateration (Fig. 3C). After the coordinate conversion, the distance from the output point and desired point can be minimized using the mean squared loss. Combining these two loss functions returns output helices compatible with the helices already placed and a midpoint along the guide. As a proof of concept, we repeated this process five times to create a helical protein backbone along the circle/line guide shown in Fig. 3D, generating 43 sets of filtered guide endpoints in 4 minutes (∼6 seconds per output). The slowest step in the process is the conversion of the distance map to coordinate space using multidimensional scaling (83% of the total time in this run) validating our approach of using backpropagation through a small model for rapid backbone generation (Table 2).

**Table 2.**

|  |  |
| --- | --- |
| Total Generated Guide Endpoints | 43 |
| Total Back-propagation Time | 42s |
| Total Time Distance Map to Point Conversion | 200s |
| Unique Looped Structures/<br>Total Structures | 24<br>702 |
| Time per Looped Unique Structure/<br>Total Structures | 52s<br>1.4s |
| Total Time | 1,246s |

#### Backbone Loop Assembly

The helices produced by the generator need three loops to complete the protein backbone. Adding loops is traditionally a computationally intensive and difficult process with the possibility of failure for each looping event, making the process inefficient. Loops that are not short or have improper backbone angles can be detrimental to a structure and result in experimental failure of the design. A fast and structurally sound method for looping is essential to make the separate helix output of the generator useful for protein design.

The generator outputs a set of helical endpoints that can be used as a guide to assemble a four-helix protein from straight helices and a loop library(Fig. 4A). The looping procedure is iterative and proceeds as follows: first, a straight helix is inserted between the first two endpoints. The first loop translates the distance between the 2^nd^ and 3^rd^ endpoints while also directing the next helix towards the 4^th^ endpoint. To find this specific loop, we took a loop library of short loops with helical adapters and measured how the loops changed the 10-long helices extended on either side via the parameterization described above. The parameters were converted to give two desired vectors and the phase: the loop translation vector from the 2^nd^ to the 3^rd^ endpoints (3 features), the helix direction vector from the 3^rd^ to the 4^th^ endpoints (3 features) and the change in helical phase (1 feature), Fig. 4B. These seven features define a kd-tree to query for a specific loop and then efficiently pull the nearest neighbors. The helical phase information is absent from the endpoint representation, but the kd-tree is queried multiple times for loops that satisfy the directional vectors with a diversity of helical phases. Sampling different helical phases attempts to produces proteins of the same topology but with different relative positioning of the side chain between helices, Fig 4C. Querying the kd-tree always returns the closest match, but it doesn’t mean that multiple phase solutions exist within the loop library. The fragment additions are not perfectly aligned to the guides, so each successive addition is queried uniquely to keep the ongoing assembly within the endpoints guides, Fig 4A. As the helix and loop fragments are assembled, they are filtered by steric clashes and deviations from the guide endpoints. The looping method was optimized towards producing a solution per endpoint set as quickly as possible. A combinatorial explosion of possible solutions can occur depending on how well the loop required by the endpoint set matches the loop library (and results in a protein without a steric clash). The number of queries checked is increased per endpoint set until a solution is found.

**Figure 4.**
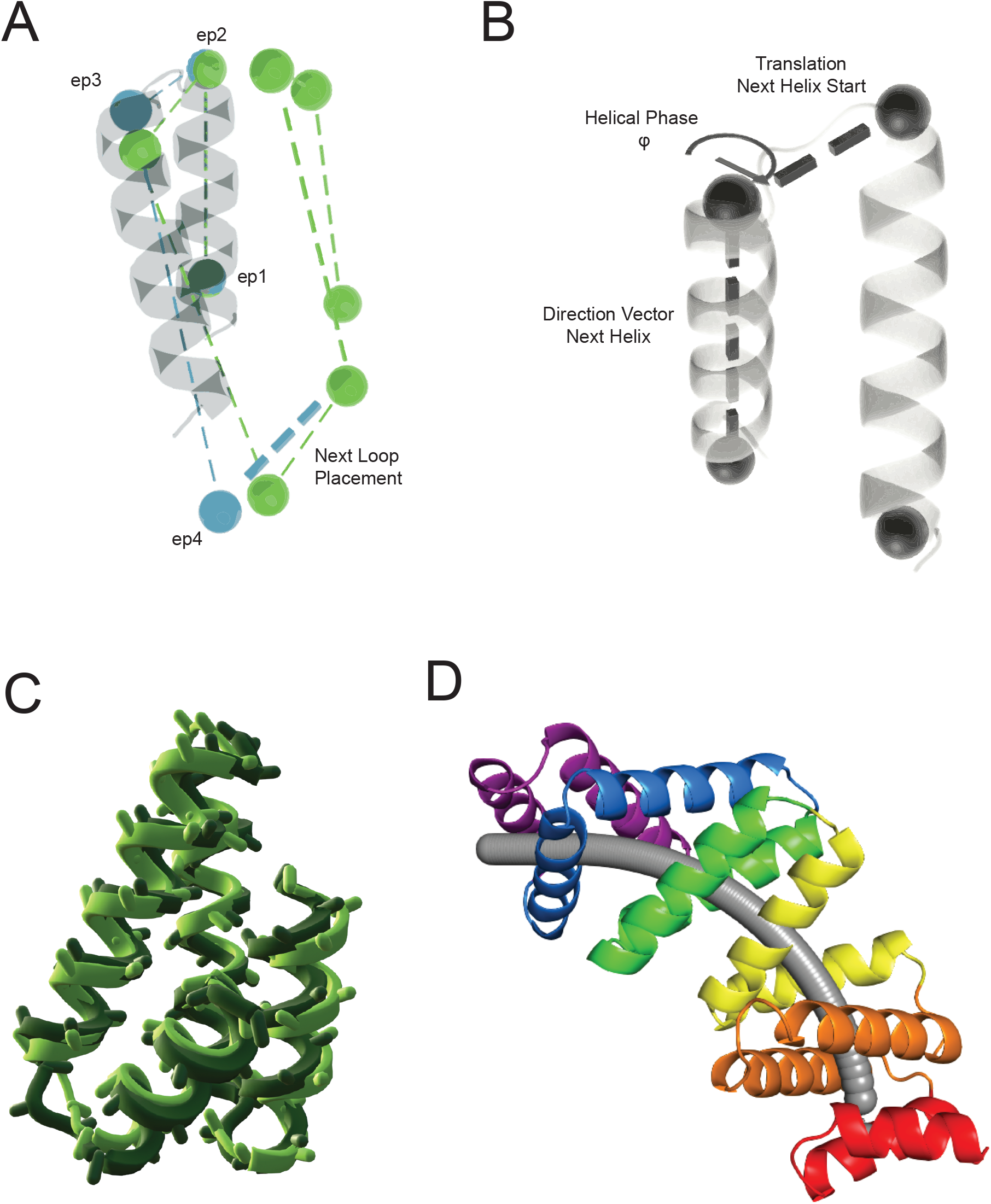
Loop Addition Protocol **(A)** Diagram for process of loop placement alternating between helix and loop addition. The guide endpoints are green while the endpoints of the actual build are blue. The next loop is queried from the current (blue) endpoint to the next guide endpoint (green). **(B)** Fitting the loop library to helical parameters. The parameterizations are converted into the translation vector between the two endpoints to be looped, the direction vector of the next helix and the helical phase. **(C)** Examples of two different loop solutions for a set of endpoints. **(D)** Looped Solution from endpoints in Figure 3D.

To test the looping protocol, the reference dataset was transformed into helical endpoints. From a random 1,028 subset of the reference dataset, 1,020 endpoint sets were successfully assembled into full proteins on average taking 0.7s per set of endpoints (Table 3). Each set of endpoints that was successfully assembled generally had many variations based on different loop selections. Loops that were queried from the same helical phase bin during assembly were counted as repeat structures to reduce the output of very similar structures. After removing the similar structures, 11,830 total backbones were produced during the assembly process (0.06 s per structure). The Gen_Full and Gen_278 generators gave similar results to the reference set, validating that the generators produce guides that can be quickly and reliably looped. This loop assembly method is not limited to the output of the generators trained in this work; any protein that can be approximated by straight helices can be fit and re-sampled using this method. We then looped the guide endpoints produced along the circle-guide from Figure 3D, (Fig. 4D). In total (back-propagation and looping), it took 52 seconds per endpoint set structure and 1.4 seconds for each structure generated in total (Table 2).

**Table 3.**

| Generator | Gen_Full | Gen_278 | Reference |
| --- | --- | --- | --- |
| Generated Examples | 1028 | 1028 | 1028<br>[Random Subset] |
| Loop Success | 1015 | 1015 | 1020 |
| Total Structures | 12,407 | 12,287 | 11,830 |
| Time Per Endpoint Set | 0.9 | 1.2 | 0.7 |
| Time Per Stucture | 0.06 | 0.10 | 0.06 |

#### Sequence Design

Rapidly produced backbones would pair well with rapid sequence design methods like a graph-based generative (GraphGen) message passing network that auto-regressively designs protein sequences more quickly and accurately than the Rosetta-based PackRotamers^7^. This network has been updated but this work was done previously though unpublished^8^. GraphGen was originally trained on crystal structures unlike the straight helix outputs of this protocol. As such, the backbones are not in the perfect location for side chain addition, like crystal structures, and backbone perturbations (>0.1 Å) will be necessary to finalize the structure. To test if the GraphGen network is robust enough to predict sequences for the straight-helix backbones, we took the reference four-helix structures and straightened the helices by constraining the C_α_ atoms of the helices to the straight helical fits and energy minimizing the poly-alanine backbone in Rosetta^37^ (Figure 5A). This forms a set of straight helical proteins with loops that have known sequences corresponding to high-quality structures with minimal structural perturbations. The GraphGen network was often confused by the straight-helix inputs as it predicted an excess of alanine residues. GraphGen predicted ∼35% of the total amino acids as alanine versus ∼5% alanine residues from the reference sequences. The alanine predictions are a logical choice of the GraphGen network from a fixed backbone perspective as alanine promotes straight helical backbones, but also poor in this context since alanine is not of sufficient hydrophobicity to promote protein folding via hydrophobic collapse in such small proteins. Final design structures were obtained by placing the predicted or reference sequences onto the straightened backbones and relaxing with Rosetta. 100 random structures were verified with a lean version of AlphaFold2 by excluding the multiple sequence alignment data^37^. The relaxed reference structures matched the AlphaFold2 predictions with a median RMSD of 0.88 Å while many of the GraphGen predictions were wildly different with a median RMSD of 8.1 Å (Figure 5B).

**Figure 5.**
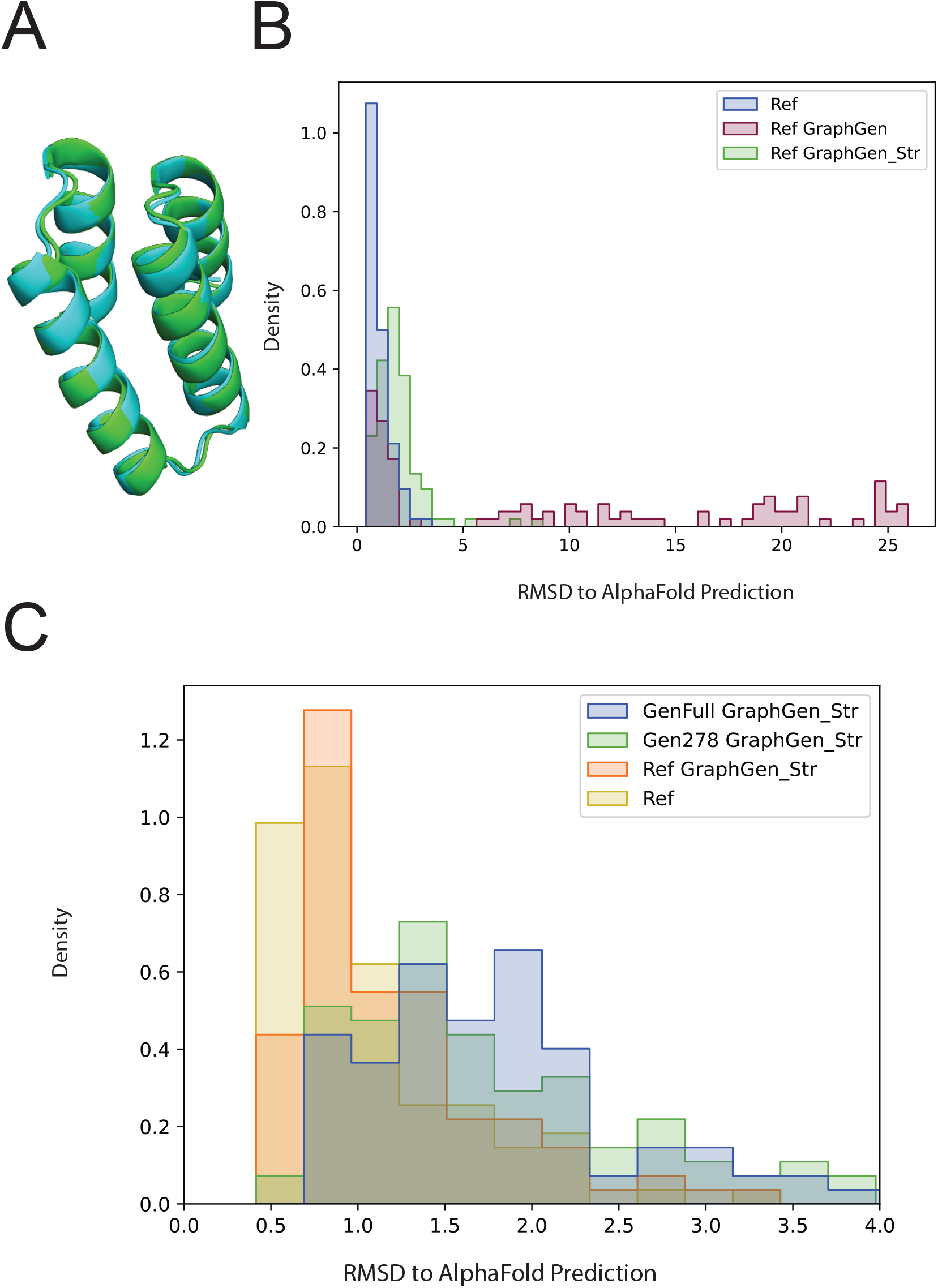
Full 4-Helix Pipeline Evaluation **(A)** Example of a straightened structure (cyan) aligned to its parent(green) for training. **(B)** Histogram of the RMSD from the AlphaFold2 prediction to a reference straightened backbone that was relaxed in Rosetta using sequences from the reference dataset, predicted by the original network^7^ (GraphGen) and predicted by the re-trained network on straightened four helix bundles (GraphGen_Str) **(C)** Histogram of the RMSD from the AlphaFold2 prediction to a helical proteins generated by Gen_Full or Gen_278, looped from the protocol in Figure 3 and sequences designed by GraphGen_Str. A single looped structure was used per endpoint set (see Methods).

We reasoned that re-training the GraphGen network using the straightened dataset as the structural input and the reference sequences as the sequence input might improve GraphGen’s sequence design performance on the output of the generator. Using the clustering from the generator validation task, we divided the straightened dataset into train and test sets on whether the terminal helices were adjacent (15,000) or apart (11,000). Accuracy metrics are for the test set are shown in Supplemental Table 1. Retraining the network on this new dataset produced GraphGen_Str which sequence predictions matched the AlphaFold2 predictions with a median RMSD of 1.0 Å (Figure 5B).

#### 4-Helix Pipeline Test

Finally, we assembled the helix generation, loop assembly and sequence design tools as a pipeline for four-helix protein design. Helical endpoints (1,028 sets) were generated from Gen_Full, diversified while adding loops and then sequence designed with GraphGen_Str, resulting in 1,015 starting topologies and 12,407 structural variants with designed sequences at a computational cost ∼0.1s per structure (∼1s per unique endpoint set). To verify the validity of these structures we took a single looped structure from 100 randomly generated endpoint sets, used GraphGen_Str to design sequence and predicted the structure with AlphaFold2 (Fig. 5C). The AlphaFold2 prediction of the sequence was compared with sequence placed on the starting backbone and relaxed with Rosetta. The pipeline produced reasonable proteins with a median RMSD to the AlphaFold2 prediction of 1.68 Å (Std. Dev 1.18 Å). Gen_278 produced similar results with a median RMSD of 1.55 Å (Std Dev: 1.46 Å). A similar analysis using Rosetta score is shown in Supplemental Figure 1.

## Discussion

Instead of a full atom model, we have developed a neural network to guide fragment assembly. Using helical parametrizations to simplify the features of a protein backbone dataset, we developed a series of neural networks and data structures to rapidly generate helical backbones. The parametrization of the helices allows the generator network to be of a small size which facilitates rapid backpropagation through the network to find specific helical arrangements. As a proof of concept, we generated a 12-helix protein that accurately reflected the desired shape. The reduced features from the fitting also allowed a similar generator to be trained using only 278 examples. Feature reduction based on secondary structure may be able to effectively train generators on many types of proteins with few examples.

Although we presented a full set of protein design methods, we imagine that incorporation/replacement of parts of the proposed pipeline with full atom models like AlphaFold2 or RFDiffusion will be beneficial for adding loops, confirming the designs or merging them with more complex structures^4,10^. Rapidly produced outputs from generators like these could help jump-start hallucination or inpainting or be used to generate secondary-structure conditioning for models like RFDiffusion.

## Python Computational Libraries

Tensorflow^38^, Pytorch^39^, Scipy^35^, Numpy^40^, Pymol^41^, Pandas^42^, Pyrosetta^43^, LMFit^44^ https://github.com/bcov77/npose, https://github.com/rdkibler/superfold_public^37^

## Hardware

All methods were run on an AMD Ryzen 7 3700X 8-Core processor (3.59 GHz) with 64 GB of available RAM or NVIDIA GeForce RTX 2070 Super.

## Code Availability

https://github.com/NickWoodall/HelixGen

## Author Contributions

R.K. and B.W created the helical fitting method. B.C. designed the reference dataset and the helical loop library as well as the methods to assemble the protein fragments (npose library). N.W. fit and clustered the dataset, trained the GAN and other machine learning methods and created the guided loop assembly method.

## Methods

### Loop Adding Fragment Assembly

The first step in the assembly process is special since the first is without a reference for phase. Essentially, the helix can be rotated around its helical axis and still be compatible with the generated guide endpoints. To query all starting phases simultaneously, we created new kd-tree from the features that was rotated around the helical axis incrementally by 10 degrees and concatenated them together to make a kd-tree 36 times larger that is only used for the first step. The rest of loop additions will be aligned to the reference, the helix-loop-helix backbone made for fitting, before querying the smaller kd-tree.

The assemblies are also optionally diversified by altering helix length rather than using the length defined by the guide endpoints which is approximate anyway. Altering the helix length drastically changes helical phase which initiates the following loop. This new starting point drastically changes the loop that can be chosen which propagates to the helical phase of the next helix resulting in significant backbone diversity. Assemblies that are sterically clashing (2.85 Angstrom cut-off) or actual endpoints that deviate more than 6 angstroms from guide endpoints were removed.

### Helical Assembly Process

The overall process is a series of two-helical additions whose midpoints follow a set of guide points. The guide points were made from a quarter circle of radius 30 angstroms using 100 points and concatenated to a line segment of 10 angstroms using 30 points that continues the circle. The start of the circle was placed at the origin for convenience with the circle/line in the positive xz-plane. To initiate the process, two random straight starting helices (2,000 total) were taken from the reference fits and placed at the origin in the xy-plane. The next target midpoint of the two helices is determined to be the point on the guide that is closest to 10 angstroms away from the current helical midpoint.

The distance map from the generator was converted to point space using trilateration in order to determine the loss to the target midpoint. Trilateration utilizes a convenient reference frame where the base-triangle of the tetrahedron is in the xy-plane at (0,0,0), (var1, 0, 0) and (var2, var3, 0) to simplify the calculation of the intersection of the three spheres. The endpoints to be buttressed are first aligned into the trilateration reference frame bringing along the ‘target’ midpoint into the loss function reference frame. Trilateration of a single point requires an assumption of ±z. The first point is assumed positive, and the remainder of the points are assigned z-values based on distances from the generated distance map. The midpoints loss is calculated as the mean squared error of the generated midpoint from the target midpoint. To maintain the two input helices, a mean squared error loss function was created between the generated distances and the input distances for the distances between the first two helical endpoints.

Each input was batched to 200 different starting points for optimization of the loss function. After 200 cycles of optimizing the loss function the outputs were filtered on loss cut-offs. Since multidimensional scaling conversions from the distance map to the output endpoints is the slow step, only a maximum of 2,000 outputs were allowed, with priority being given to outputs from different endpoint sets. The outputs were tested for clashes (of straight helices interpolated between endpoints) and the final output midpoint was allowed to deviate by 5 angstroms. The final output midpoints were moved back into the original reference frame using the Kabsch algorithm to align the four input endpoints to first four generated endpoints. The process is repeated until there are no guide endpoints 10 angstroms away.

#### AlphaFold2 Verification of the 4-Helix Pipeline

Distance maps were generated from random inputs that were then converted to helical endpoints sets. Loops were adding using the loop-fragment assembly process described above. To save computational time, one structure from each helical endpoint sets was chosen for structure prediction by AlphaFold2. The backbone with the highest percent core of residues calculated the based on C_α_/C_β_ atom vector overlap was chosen to avoid choosing a particularly bad backbone, but it is likely uncorrelated to the “best” backbone. The predicted sequence was placed on the chosen backbone and relaxed in Rosetta for comparison the AlphaFold2 predictions. The AlphaFold2 predictions were run on a reduced network that doesn’t utilize the multiple sequence alignments which is used to quickly evaluated simple proteins like these. The prediction was run on 5 different model parameters sets and the lowest RMSD prediction to the relaxed structure was used. RMSD measurements were kept regardless pLDDT scores since removing unsure predictions decreased the median RMSD and likely indicated a poor structure which is reflected in the RMSD from design to AlphaFold prediction.

## Supplemental Figure Legend

**Supplemental Figure 1.**
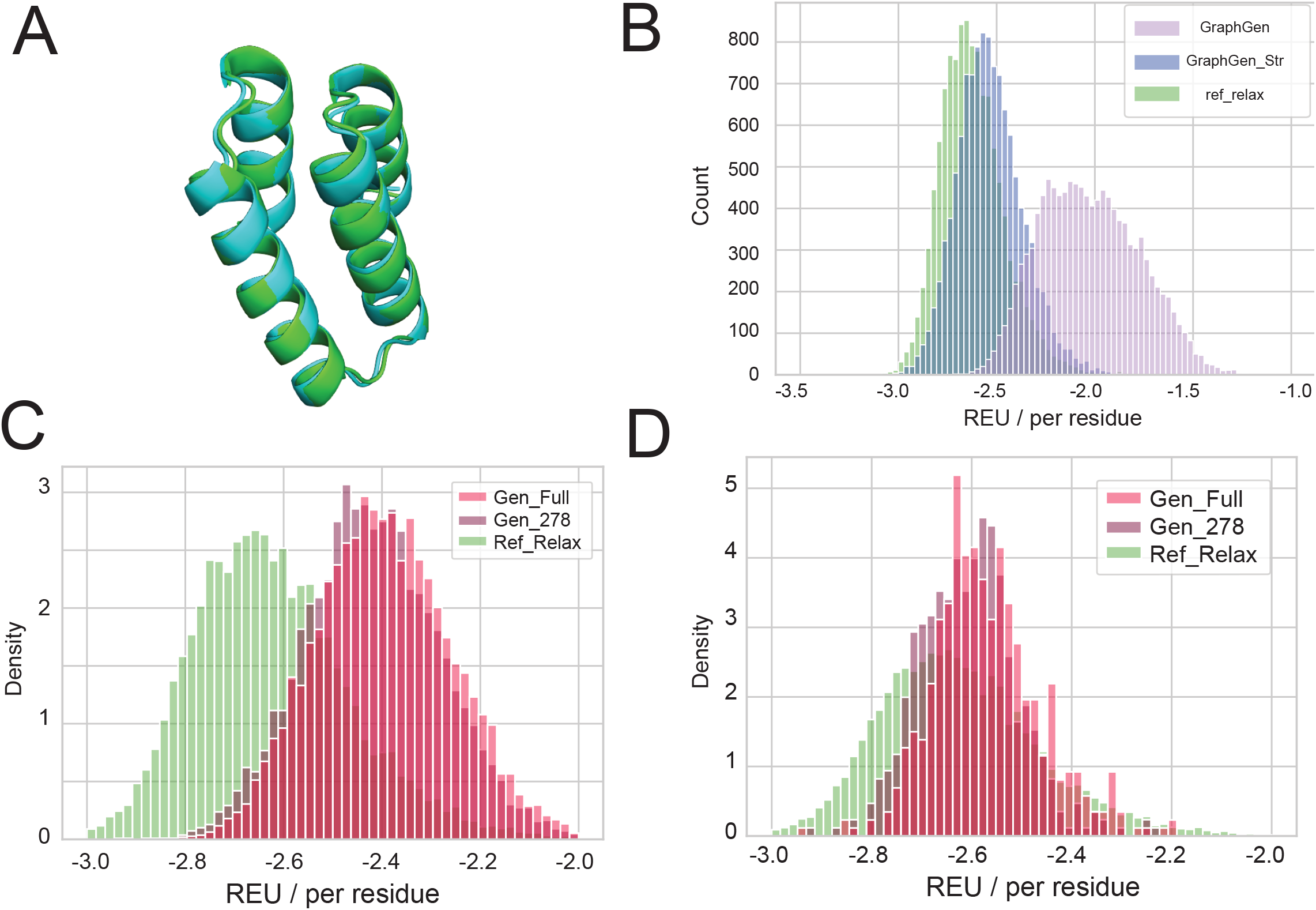
Sequence design with GraphGen **(A)** Example of a straightened structure (cyan) aligned to its parent(green) for training. **(B)** Histogram of the scores in Rosetta Energy Units/ residue for the straightened reference set relaxed with the reference sequences (ref_relax), sequences predicted by the original network^7^ (GraphGen) and sequence predicted by the re-trained network on straightened four helix bundles (GraphGen_Str) **(C)** Rosetta scores in REU/res for proteins with generated and looped backbones, sequences predicted by GraphGen_Str and then relaxed in Rosetta. **(D)** Best scoring selections from each set of endpoints in part (C).

**Supplemental Table 1.**

|  | Model Accuracy |  |
| --- | --- | --- |
|  | GraphGen | GraphGen_Str |
| Hydrophillic Group<br>D, E, H, K, N, Q, R, S, T | 58% | 95% |
| Hydrophobic Group<br>A, I, L, V, F | 81% | 96% |
| A, F, I, L, V | 75%, 9%, 9%, 35%, 8% | 74%, 27%, 43%, 81%, 59% |
| Glycine, Proline | 15%, 4% | 93%, 96% |

## References

1. Wang, J. et al. Scaffolding protein functional sites using deep learning. (2022).

2. Wicky, B. I. M. et al. Hallucinating symmetric protein assemblies. Science 378, 56–61 (2022).

3. Lutz, I. D. et al. Top-down design of protein nanomaterials with reinforcement learning. http://biorxiv.org/lookup/doi/10.1101/2022.09.25.509419 (2022) xdoi:10.1101/2022.09.25.509419.

4. Watson, J. L. et al. Broadly applicable and accurate protein design by integrating structure prediction networks and diffusion generative models. http://biorxiv.org/lookup/doi/10.1101/2022.12.09.519842 (2022) xdoi:10.1101/2022.12.09.519842.

5. Lee, J. S., Kim, J. & Kim, P. M. ProteinSGM: Score-based generative modeling for de novo protein design.

6. Norn, C. et al. Protein sequence design by conformational landscape optimization. Proc. Natl. Acad. Sci. 118, e2017228118 (2021).

7. Ingraham, J., Garg, V., Barzilay, R. & Jaakkola, T. Generative Models for Graph-Based Protein Design.

8. Dauparas, J. et al. Robust deep learning–based protein sequence design using ProteinMPNN. Science 378, 49–56 (2022).

9. Mirdita, M. et al. ColabFold: making protein folding accessible to all. Nat. Methods 19, 679–682 (2022).

10. Jumper, J. et al. Highly accurate protein structure prediction with AlphaFold. Nature 596, 583–589 (2021).

11. Anishchenko, I. et al. De novo protein design by deep network hallucination. Nature 600, 547–552 (2021).

12. Baek, M. & Baker, D. Deep learning and protein structure modeling. Nat. Methods 19, 13–14 (2022).

13. Anand, N. & Huang, P. Generative modeling for protein structures.

14. Eguchi, R. R., Choe, C. A. & Huang, P.-S. Ig-VAE: Generative Modeling of Protein Structure by Direct 3D Coordinate Generation.

15. Sabban, S. & Markovsky, M. RamaNet: Computational de novo helical protein backbone design using a long short-term memory generative neural network.

16. Chen, Z. et al. De novo design of protein logic gates. Science 368, 78–84 (2020).

17. Woodall, N. B. et al. De novo design of tyrosine and serine kinase-driven protein switches. Nat. Struct. Mol. Biol. 28, 762–770 (2021).

18. Silva, D.-A. et al. De novo design of potent and selective mimics of IL-2 and IL-15. Nature 565, 186–191 (2019).

19. Langan, R. A. et al. De novo design of bioactive protein switches. Nature 572, 205–210 (2019).

20. Foight, G. W. et al. Multi-input chemical control of protein dimerization for programming graded cellular responses. Nat. Biotechnol. 37, 1209–1216 (2019).

21. Huang, P.-S. et al. High thermodynamic stability of parametrically designed helical bundles. Science 346, 481–485 (2014).

22. Ng, A. H. et al. Modular and tunable biological feedback control using a de novo protein switch. Nature 572, 265–269 (2019).

23. Sahtoe, D. D. et al. Reconfigurable asymmetric protein assemblies through implicit negative design. Science 375, eabj7662 (2022).

24. Courbet, A. et al. Computational design of mechanically coupled axle-rotor protein assemblies. (2022).

25. Hsia, Y. et al. Design of multi-scale protein complexes by hierarchical building block fusion. Nat. Commun. 12, 2294 (2021).

26. Ben-Sasson, A. J. et al. Design of biologically active binary protein 2D materials. Nature 589, 468–473 (2021).

27. Yeh, C.-T., Obendorf, L. & Parmeggiani, F. Elfin UI: A Graphical Interface for Protein Design With Modular Building Blocks. Front. Bioeng. Biotechnol. 8, 568318 (2020).

28. Huang, P.-S. et al. RosettaRemodel: A Generalized Framework for Flexible Backbone Protein Design. PLoS ONE 6, e24109 (2011).

29. Guffy, S. L., Teets, F. D., Langlois, M. I. & Kuhlman, B. Protocols for Requirement-Driven Protein Design in the Rosetta Modeling Program. J. Chem. Inf. Model. 58, 895–901 (2018).

30. Correia, B. E. et al. Proof of principle for epitope-focused vaccine design. Nature 507, 201–206 (2014).

31. Yang, C. et al. Bottom-up de novo design of functional proteins with complex structural features. Nat. Chem. Biol. 17, 492–500 (2021).

32. Grigoryan, G. & DeGrado, W. F. Probing Designability via a Generalized Model of Helical Bundle Geometry. J. Mol. Biol. 405, 1079–1100 (2011).

33. Goodfellow, I. J. et al. Generative Adversarial Networks. Preprint at http://arxiv.org/abs/1406.2661 (2014).

34. Pedregosa, F. et al. Scikit-learn: Machine Learning in Python. Mach. Learn. PYTHON.

35. Virtanen, P. et al. SciPy 1.0: fundamental algorithms for scientific computing in Python. Nat. Methods 17, 261–272 (2020).

36. Shaham, U. et al. SpectralNet: Spectral Clustering using Deep Neural Networks. Preprint at http://arxiv.org/abs/1801.01587 (2018).

37. Kibler, R., Broerman, A. & Leung, P. SuperFolder Public.

